# Smart switching in feedforward control of grip force during manipulation of elastic objects

**DOI:** 10.1101/194407

**Authors:** Olivier White, Amir Karniel, Raz Leib, Charalambos Papaxanthis, Marie Barbiero, ilana Nisky

## Abstract

Switching systems are common in artificial control systems. Here, we suggest that the brain adopts a switched feedforward control of grip forces during manipulation of objects. We measured how participants modulated grip force when interacting with soft and rigid virtual springs when stiffness varied nearly continuously between trials. We identified a sudden phase transition between two forms of feedforward control that differed in the timing of the synchronization between the anticipated load force and the applied grip force. The switch occurred several trials after a threshold stiffness level. These results suggest that in the control of grip force, the brain acts as a switching control system. This opens new research questions as to the nature of the discrete state variables that drive the switching.

## Introduction

A driver switches between different gears, air conditioners switch between on and off states and irrigation mechanisms switch between closed and open circuits. In control theory, hybrid systems are systems with continuous and discrete states. The examples outlined above are switched systems, a subclass of hybrid systems, that are defined as continuous time systems with isolated discrete switching events (Liberzon, 2003). The discrete switching often occurs based on a threshold value of another continuous variable, e.g. the velocity in the former example, the temperature of the thermostat in the second and moisture in the last. Such control systems have many benefits, including economy in control effort (Ben-Itzhak and Karniel, 2008; Karniel, 2011; Leib and Karniel, 2012) and the ability to stabilize otherwise unstable systems (Margaliot and Liberzon, 2006; Lin and Antsaklis, 2009; Liberzon, 2003; Wicks et al., 1998).

Switching is also common in human control of movement. For example, human movements are intermittent (Craik, 1947; Navas and Stark, 1968; Neilson et al., 1988; Miall et al., 1993; Doeringer and Hogan, 1998; Squeri et al., 2010; Gawthrop et al., 2014), they switch between different types (Kelso, 1984; Levy-Tzedek et al., 2011, 2010) and neural activity states (Gross et al., 2002; Loewenstein et al., 2005; Yartsev et al., 2009). Several models based on switching were proposed to describe control of standing (Bottaro et al., 2005; Asai et al., 2009; Gawthrop et al., 2014), stick balancing (Gawthrop et al., 2013), and hand movements (Ben-Itzhak and Karniel, 2008; Leib and Karniel, 2012). Intermittent control was proposed to be at least as efficient as continuous control (Loram et al., 2011). Here we present evidence suggesting that the feedforward control of grip force during object manipulation is a switched control system, and we mention several candidate variables that correlate with the switching and that are therefore worth exploring in future investigations.

Many studies have used the modulation of grip force with anticipated load force as an evidence for prediction in the control of voluntary movement (Johansson and Westling, 1984, 1988). Moving an object held in precision grip requires the anticipation of inertial and gravitational forces that may cause its slippage (Flanagan et al., 1993; Flanagan and Wing, 1995). The anticipatory adjustment of grip force generalizes to less usual forms of load force including those dependent on object position (Descoins et al., 2006; Danion and Sarlegna, 2007; Leib et al., 2015; Sarlegna et al., 2010), velocity (Flanagan et al., 2003; Nowak et al., 2004), modified gravitational forces (Augurelle et al., 2003; White, 2015) and when forces are generated by whole body actions such as walking or jumping (Gysin et al., 2008). Without exception, when load forces are generated by a direct action of the body on the environment, grip force and load force profiles match closely as usually quantified by close-to-zero lags between their peak values, or close-to-zero lags in peak cross-correlation between them.

These studies present evidence for the anticipation of smoothly varying, often self-generated, forces. However, in many natural object manipulation tasks, the central nervous system must also adjust grip forces to deal with impulse-like destabilizing forces induced by the nearly instantaneous contact between an object and a hard surface. Several studies also addressed the control of grip force in impact-like tasks: When participants had to anticipate a sudden increase of weight after dropping a ball in a hand-held receptacle (Johansson and Westling, 1988; Bleyenheuft et al., 2009), when opening a drawer to its mechanical stop (Serrien et al., 1999), when hitting an object against a pendulum (Turrell et al., 1999) or a surface (White et al., 2011, 2012) or in a step-down task (Ebner-karestinos et al., 2016). A common observation was the occurrence of a maximum of grip force approximately 60 ms after peak load force that signed the impact. A natural question occurred as to whether this delayed grip force peak resulted from a feedback process. Recently, by studying grip force in catch trials, where load forces are not applied, experiments unambiguously demonstrated this behaviour reflects a feedforward process and is not a mere reflex response to a perturbation signal (Bleyenheuft et al., 2009; White et al., 2011). Nonetheless, this feedforward strategy contrasts sharply with the zero-delay coupling observed between grip and load forces when the latter vary smoothly. To sum up, past investigations showed that grip force control in soft and stiff elastic dynamics exhibits different feedforward control strategies. This is surprising since the underlying mechanics is described by a single stiffness parameter (k) that varies continuously.

Here, we set out to explore the nature of the transition between these two different feedforward control strategies. We studied grip force adjustment during repeated interactions with virtual objects rendered as elastic force fields. In the repeated interactions, the objects properties varied between soft objects to rigid surfaces or vice versa, resulting in systematically changing impact forces, either increasing or decreasing. We hypothesized that if participants adopt a continuous control strategy, when the stiffness will increase (or decrease) continuously over trials, the grip force – load force delay will continuously increase (or decrease) with respect to the impact. Alternatively, if participants adopt a switching control strategy, we expect to find a stiffness level around which there will be a phase transition in the synchronization between the modulation of grip force and the anticipated load force.

## Materials and Methods

### Participants

Eighteen right-handed adults (14 females, 20 to 40 years old, mean=24.3, SD=10.2 years) participated voluntarily in the experiment. All participants were healthy, without neuromuscular disease and with normal or corrected to normal vision. The experimental protocol was carried out in accordance with the Declaration of Helsinki (1964), the procedures were approved by the local ethics committee of Université de Bourgogne and a written informed consent was obtained from all participants. All participants were naïve as to the purpose of the experiments and were debriefed after the experimental session.

### Apparatus and stimuli

Participants sat in front of a virtual haptic environment with their head on a chin rest. A mini40 force-torque sensor (ATI Industrial Automation, NC, USA) was mounted on the handle of a robotic device (Phantom 3.0, Sensable Technologies, RI, USA) to record grip force (normal force component, -Fz). The 3d positions and forces of the robotic arm were controlled in closed loop at 1kHz. Participants looked into two mirrors that were mounted at 90 degrees to each other, such that they viewed one LCD screen with the right eye and one LCD screen with the left eye. This stereo display was calibrated such that the physical location of the robotic arm was consistent with the visual disparity information.

### Experimental procedure

Participants grasped the force sensor with a precision grip (thumb on one side and index finger on the other side, Fig. 1A inset). To initiate a trial, participants moved their right hand, displayed as a grey 5-mm sphere, into another grey starting sphere (10mm diameter), displayed at body midline and at chest height. Then, a green target rectangle (12cm width, 10mm height) appeared 15cm above home position (Fig. 1A). Participants were instructed to move the cursor straight upward to touch the target and bring it immediately back to home position without stopping at the reversal point. No instructions were provided regarding how they had to adjust grip force. To avoid large trial-to-trial variability in movement kinematics, after each trial, a line was displayed at a height proportional to peak velocity together with lower (45cm/s) and upper (55cm/s) bounds. The color of the line was red if peak velocity was outside the interval or green in successful trials (Fig. 1A). Participants adjusted their movement such that peak velocity fell in that interval.

**Figure 1.**
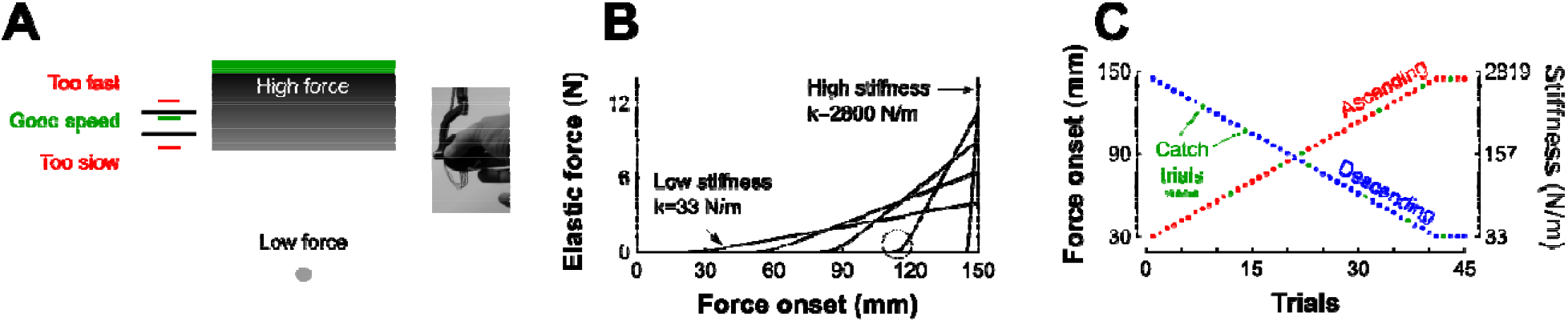
Description of experimental procedures. (A) A grey sphere was moved toward a green target through a parameterized elastic force field. The grey gradient represents the magnitude of the force field for a given stiffness. It is low far from the target (bright grey) and increases when vertical position approaches the target (dark grey). Left: feedback of achieved peak velocity. The right inset depicts the force sensor attached to the end of the robot handle held in precision grip. (B) Examples of force-position trajectories of five elastic force fields parametrized by five pairs of stiffness levels and force onset position. Force onset occurred between 3cm and 14.5cm above home position. The steeper the slope the stiffer the force field. The circle highlights the second order interpolation between a null force field and a linear elastic force field. (C) Structure of “Ascending” and “Descending” blocks, where stiffness increases (red) and decreases (blue), respectively. Green dots represent catch trials, for which the stiffness is set to 0 N/m.

The target was located inside an elastic force field F that was haptically rendered according to 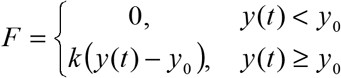, where yO is the boundary of the object and k the stiffness value. Such force field emulates a one-sided spring-like object that only resists compression. The more the cursor approached the target, the more effort was required to move it to the target (see grey gradient in Fig. 1A). The stiffness of the force field and its onset were varied systematically such that force fields with higher levels of stiffness also onset further along the movement progression (as depicted in Fig. 1B for five different force fields). The weakest elastic force field was generated when force onset occurred at 3cm and linearly ramped up to a maximum force of 4N along the 12 remaining cm (Fig. 1B, k=4/0.12=33N/m). Similarly, the strongest force field was obtained when force onset occurred 5mm below the target’s lower surface and ramped up to a maximum force of 14N (Fig. 1B, k=14/0.005=2800N/m). Force onset (yO) and stiffness (k) pairs, were parameterized independently trial by trial. The transitions between zero force outside of the elastic force field and non-zero force was smoothed with a second order polynomial interpolation (Fig. 1B, circle) to avoid mechanical vibrations and overheating of the robot motors, particularly in stiff trials. Consequently, movements were felt as natural and continuous.

The recording session consisted of 10 blocks with 45 movements in each block. In the first five blocks, the stiffness of the elastic force field increased during 41 trials and plateaued for the last 4 trials (Fig. 1C, red “Ascending” blocks). Force onsets were linearly spaced between 3cm and 14.5cm by steps of exactly 0.2875cm. In the last five blocks, force field stiffness decreased over trials (Fig. 1C, blue “Descending” blocks). Eight participants started the experiment with the “Ascending” blocks and 7 participants started the experiment with the “Descending” blocks. In every block, 6 trials (13%) were randomly chosen to be catch trials in which the stiffness and onset of the elastic force field were set to zero, effectively vanishing the force field (Fig. 1C, green disks). Their order was counterbalanced between participants. In the remaining 39 trials, the natural dynamics remained intact.

### Data processing and statistical analyses

Position and grip forces were recorded at 500Hz. Grip force rate, velocity and acceleration were obtained using a central-difference algorithm and smoothed with a zero phase-lag autoregressive filter (cutoff 20Hz). All trials were aligned to movement onset, defined as the time when velocity went above 3cm/s during at least 100ms. We also recorded temporal occurrences and values of peak acceleration and grip forces (minmax function in matlab and visual check). Finally, we extracted the value of grip force rate when the hand reached a height of 3cm, or, put differently, at expected force onset. This measure provides an estimate of feedforward mechanisms of grip force and allows direct comparison between normal and catch trials. To compare real and sparse catch trials, we grouped trials of the same block in 7 mini blocks of 5 trials each (except the first and last mini blocks with 10 trials each). That way, every mini block had both real and catch trials.

We verified that starting with five “Ascending” blocks (N=10) or five “Descending” blocks did not influence any of the above variables (all F_1,14_<1.3, p>0.324). We therefore pooled these two groups together. Quantile-quantile plots were used to assess normality of the data. A three-way ANOVA was conducted on the above variables to assess the effects of stiffness (Mini block, 1 to 7), Block condition (“Ascending” vs. “Descending”) and Type of trial (“Real” vs. “Catch”). Paired t-tests of individual subject means or bootstrap procedures were used to investigate differences between conditions on the above variables. Significance level was set to alpha=0.05. Data processing and statistical analyses were done using Matlab (The Mathworks, Chicago, IL). Linear fits were calculated with the polyfit function. Partial eta-squared values are reported for significant results to provide indication on effect sizes.

## Results

Participants grasped a force transducer attached to the handle of a haptic device and produced vertical arm movements to touch a virtual target situated 15cm above home position (Fig. 1A). The robotic device generated a resistive vertical elastic force field that was parameterized by the stiffness of the field. As trials progressed, the stiffness of the force field either increased or decreased between two extremes, and force onset was shifted further from or closer to force onset, depending on block condition (“Ascending” and “Descending”, respectively, Fig. 1C). Force fields with the lowest stiffness were similar to a soft elastic force fields that are typically used in other studies (Descoins et al., 2006). In contrast, the force fields with the highest stiffness resembled collisions between the hand-held device and a rigid surface (White et al., 2011, 2012). To measure the feedforward grip force adjustment, we interspersed catch trials, in which visual information was available to the participants but no forces were applied. We explored the transition in grip force control between these two extremes.

## Grip force is different when interacting with high-impact and low-impact elastic force fields

Figure 2 illustrates high-stiffness (left column, solid line) and low-stiffness (right column, solid line) trials averaged across blocks and participants in a force field trial (black line) and in a catch trial (green line). In the high-stiffness condition, the vertical position increased until the target was touched at 15cm and then decreased to return to the home position. Participants achieved mean peak velocities of 49.9cm/s (SD=8.3cm/s), within the prescribed 45-55cm/s interval. The elastic force field was null until the position of the hand reached the boundary of the field at x=14.5cm and then increased up to 13.75N (middle row, black line). The vertical pushing force increased when the cursor approached the target and decreased on its way back to the starting position. Grip force increased first to counteract the inertial force (Fig. 2, Acceleration row) induced by accelerating the mass of the device and exhibited a first local peak synchronized with a local peak in the load force. Then, after a small dip, grip force increased again in anticipation of the mechanical event. This is also reflected by positive grip force rates for 200ms before impact (bottom row). However, in this stiffness condition, peak grip force was clearly delayed by 40 ms (SD=6ms) after the peak of the elastic force field.

**Figure 2.**
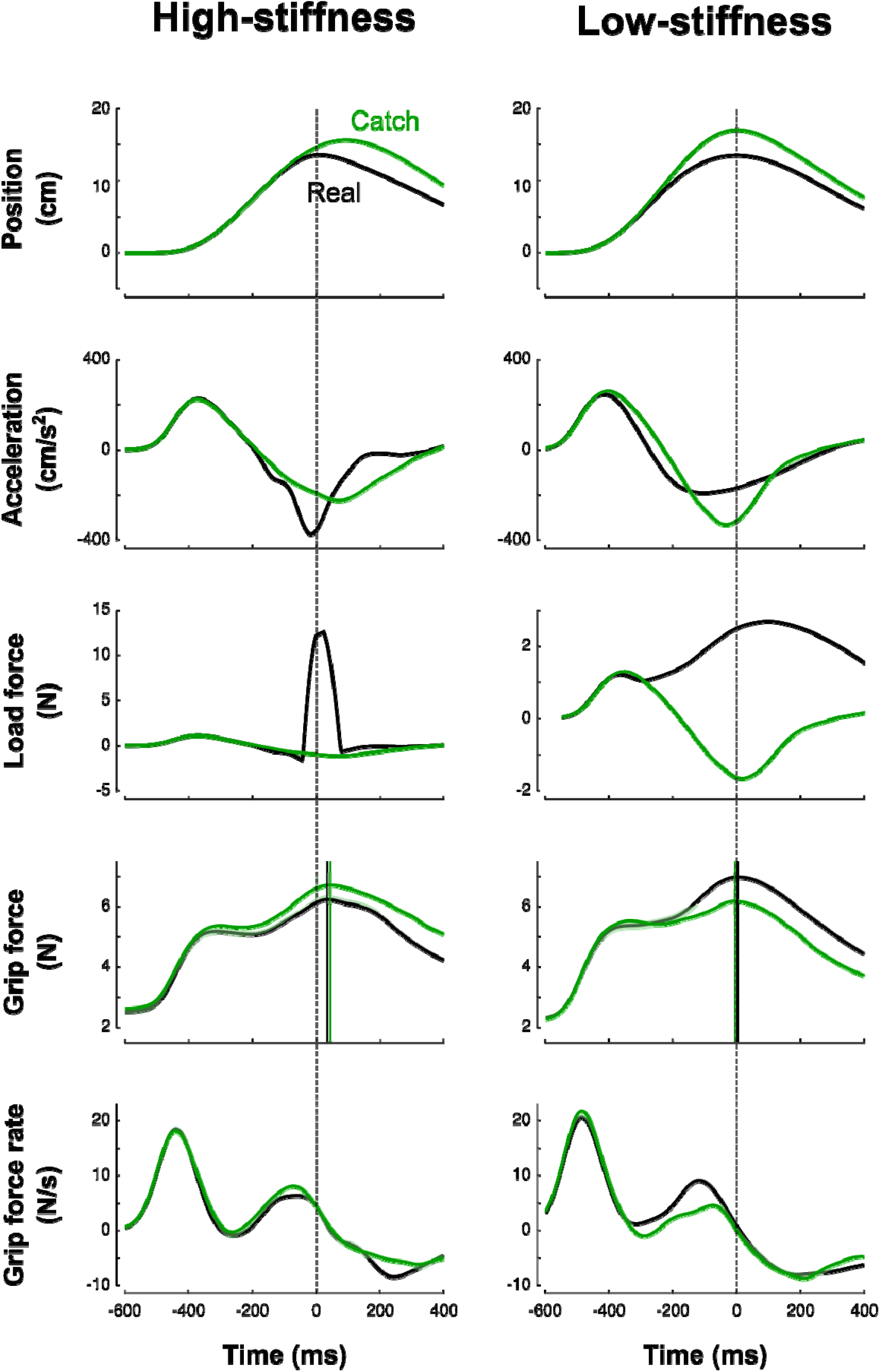
Averaged traces corresponding to high-stiffness (left column) trials and low-stiffness (right column) trials across blocks and participants. Top to bottom: vertical position, vertical acceleration, load force, grip force and grip force rate are depicted as a function of time. Black and green lines correspond to real and catch trials, respectively. All traces are aligned with peak of impact (vertical dashed cursor across panels, time 0). Cursors for the grip force traces are positioned at their respective maximum. The lags calculated between peaks of grip and elastic forces illustrate the difference between high-stiffness (lag=40ms, SD=6ms) and low-stiffness (lag=4ms, SD=6ms) conditions. Error shade areas correspond to SEM. Traces are not normalized.

In the low-stiffness trials (Fig. 2, right column), the position and acceleration trajectories resembled those for the high-stiffness trials. However, the elastic force field was smoother: it increased for 400ms up and reached a 4-N peak. Grip force and grip force rates paralleled the traces observed in the stiff condition with one notable difference: peak grip forces were synchronized with the impact, both in real and catch trials (mean=4ms, SD=6ms).

## Motor planning is similar between real and catch trials

It is important to stress that the delay between grip force peak and load force peak observed in high-stiffness trials is not a consequence of a feedback control, but rather a feedforward control that includes a delay. We observed the same behavior in catch trials without the presence of the resistive elastic forces (Fig. 2, green lines). The green vertical line is positioned at the time when peak elastic force would have occurred given the vertical position. In particular, grip force peaks were delayed by a similar amount relative to impact or expected impact in both real and catch trials.

Due to the absence of resistive forces, different kinematic profiles after t=0 (the anticipated onset of the perturbation) were induced, and the peak position overshot the target. Moreover, in these trials, participants expected a force ramp but did not feel it. It could be argued that these errors signals could have driven a feedback adjustment of grip force rather than reflect a feedforward strategy even in catch trials. We conducted complementary analyses to show that several parameters characterizing motor planning are the same between real and catch trials.

First, we extracted the value of grip force rate, a reliable index of feedforward grip force control (White et al., 2008), at the time of expected force rise. A t-test failed to report a difference between real and catch trials on grip force rate (t_17_=0.5, p=0.642).

Second, we examined the trial-by-trial variations, and verified that grip force rate in catch trials were not statistically different from grip force rate in the real trial that immediately preceded or succeeded them. To do so, we defined two additional variables by subtracting grip force rate in the previous (R_t−1_) or next real trial (R_t+1_) from grip force rate in the catch trial (C_t_) between them (C_t_-R_t−1_ and C_t_R_t+1_). The ANOVA reported no difference for C_t_-R_t−1_ (Mini block: F_6,238_=1.8, p=0.109; Block condition: F_1,238_=0.7, p=0.414) and for C_t_-R_t+1_ (Mini block: F_6,238_=0.5, p=0.821; Block condition: F_1,238_=1.0, p=0.329).

Third, to further support our claim that grip force is planned and executed using a feedforward controller, we made scatter plots between load forces and grip forces between movement onset and 100ms before peak load force (Leib et al., 2015; White, 2015; Flanagan and Wing, 1995). We then calculated, for each trial, the linear regression between these two variables and compared the offset and slopes between real and catch trials. We averaged these variables per participant and conducted an ANOVA on offsets and slopes (Catch vs. Real trial). The analysis failed to report any significant difference between catch and real trials on offsets (F_1,194_=0, p=0.969) and slopes (F_1,194_=1.0, p=0.319).

Finally, we conducted a last analysis to show that acceleration profiles over time were not statistically different between real and catch trials up to a certain time point. Indeed, Figure 2 shows that acceleration traces diverge around the perturbation induced by the elastic force field. To do so, we considered mini blocks because they included real and catch trials. We averaged acceleration traces in real trials and in catch trials separately, per participant and per mini block. Then, we ran an independent iterative t-test that compared values of acceleration between both trial types, from trial onset to maximum elastic force (time=0), and by 20-ms bins. This allowed us to extract the exact time point from which both acceleration traces diverged significantly during at least 150ms (p<0.05). We identified a divergent point unambiguously on every averaged acceleration profile in mini blocks (all t_17_>4.1, all p<0.001, all 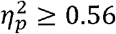). In other words, acceleration values between catch and real trials exhibited a divergent point some 30 ms after force onset in every stiffness condition. This analysis clearly shows that motor planning, in terms of its consequences measured through acceleration, is not affected by trial type.

Based on these analyses, we conclude that in both trial types, (1) motor planning was similar and (2) grip forces that lagged the load force were feedforward.

## Grip force switches between different control strategies

At the individual trial level, grip force always exhibited a clear peak over time (see Fig. 2). Interestingly, the distribution of grip force peaks themselves as a function of stiffness of the elastic force field also reached a global extremum, both in “Ascending” and “Descending” blocks (Fig. 3A, average across participants). This observation also held in individual subjects except for subject 14 in the “Descending” block condition (Fig. 3B). To compare these global grip force extrema as a function of stiffness, we fitted polynomial models to the data averaged across participants (Fig. 3A, dashed lines) or for each participant (Fig. 3B, dashed lines). Since inter-subject variability was large, we adopted a bootstrap method to test whether grip force peaks of subject fits occurred at the same stiffness level between “Descending” and “Ascending” block conditions. Zero difference between both population means was outside the 95%-confidence interval (0.08 to 0.77). Hence, we cannot reject the null hypothesis and conclude that the extremum for the “Descending” block condition occurred for lower stiffness than in the “Ascending” block condition.

**Figure 3.**
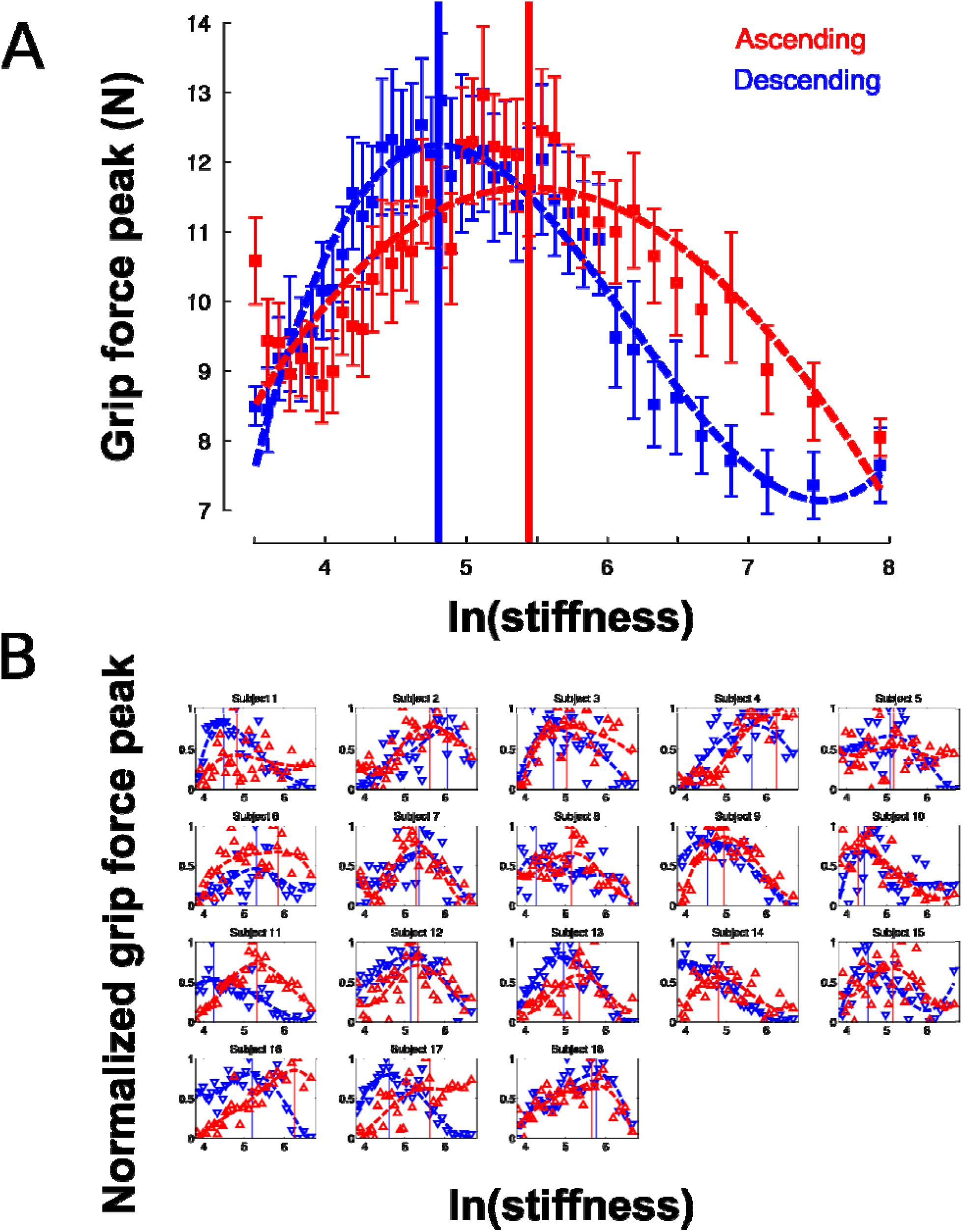
Grip force is adapted to stiffness. (A) Distribution of grip force maximum (averaged across participants) in function of ln(stiffness) in “Ascending” (red trace) and “Descending” block conditions (blue trace). The dashed lines are the best fourth order polynomial fits of each series. Goodness of fit: r^2^=0.72 in “Ascending” condition and r^2^=0.84 in “Descending” condition. The vertical cursors are positioned at peak grip force (“Ascending”: 5.07; “Descending”: 4.89, which correspond to stiffness of 159.7N/m and 133.2N/m, respectively). Error bars are between participants SE. (B) Individual normalized plots showing the same behaviour at the subject level (except for subject 14 in the “Descending” condition).

Another strategic change in grip force control is illustrated in Figure 4. The difference between times of grip force peaks and times of elastic force peaks is depicted as a function of the natural logarithm of stiffness (no significant difference between block conditions, t_17_=0.04, p=0.973), for all subjects together (Fig. 4A) or individually (Fig. 4B). The individual plots in Figure 4B reveal that most of the participants had a prominent transition in the dependency of the time difference as a function of natural logarithm of stiffness around a threshold value. Indeed, the delay seems to linearly increase from negative (leading latencies) and then plateau to a positive value (lagging time).

**Figure 4.**
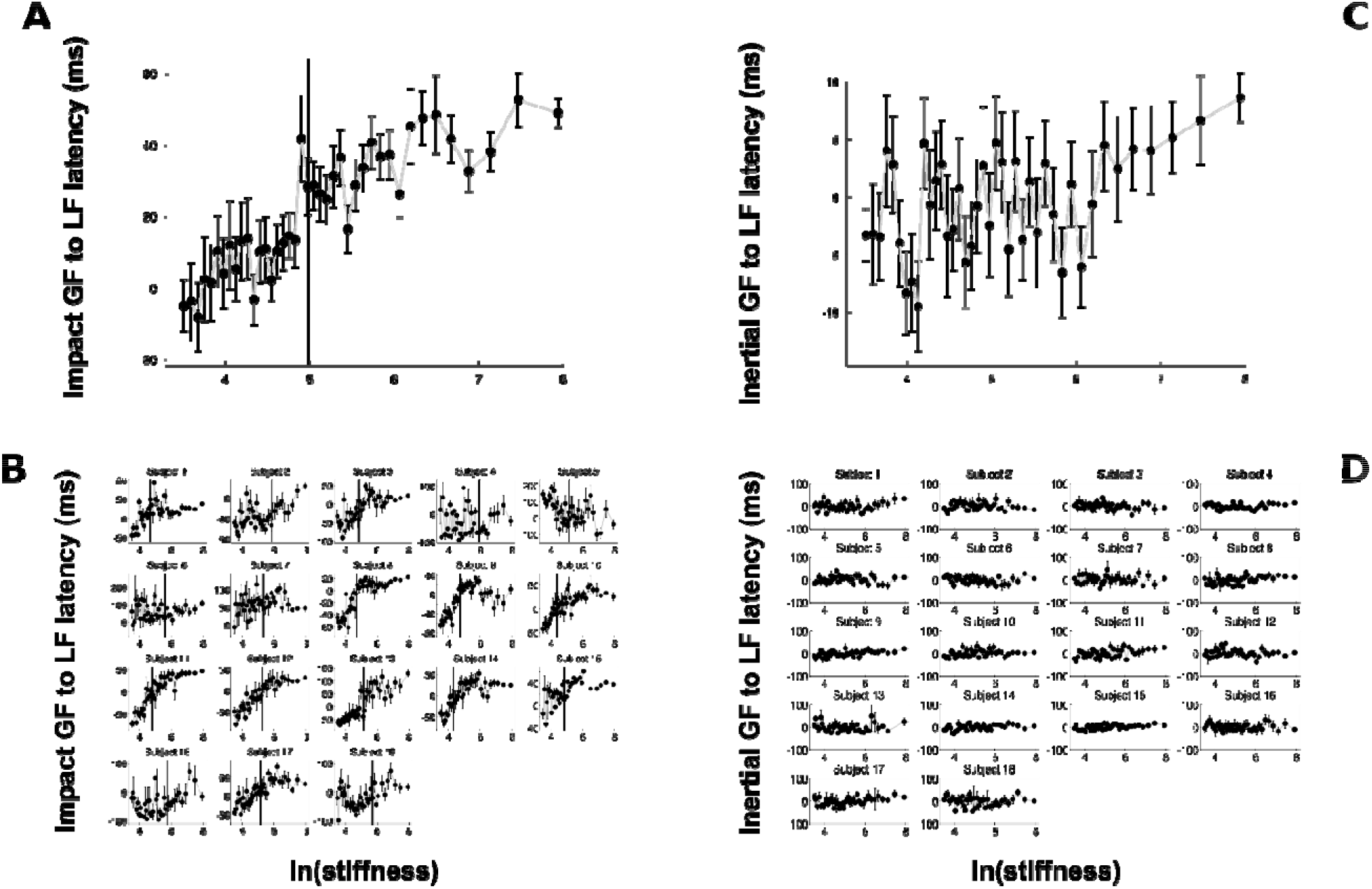
Grip force control switches for certain stiffness values. (A) Average lags across participants between grip force peak and impact (A, *Impact*) and average lags across participants between grip force and load peaks during the inertial phase of the trial (C, *Inertia*). The same lags are plotted for individual participants in the lower panels (B, Impact and D, Inertial). The lags vary with increasing stiffness during the impact phase in (A), but are constant and zero during the inertial stage (C). All variables are plotted as a function of the natural logarithm of stiffness. Positive values of latencies mean that grip force peaks lag load force peaks. Black cursors are positioned at the ln(stiffness) identified from grip force peaks that correspond to the average between the “Ascending” and “Descending” conditions. Error bars are SEM.

To reliably quantify this effect, we first extracted slopes of the linear fits of the lag in function of ln(stiffness) on individual subject data. We partitioned the data in a low-stiffness and high-stiffness subset, according to the individual thresholds (averaged between the two block conditions) estimated as the value ln(stiffness) for which peak grip force occurred (see Fig. 3B). We then statistically tested whether slopes differed between both stiffness conditions. To do so, we defined a random variable as the absolute difference of slopes between low and high stiffness conditions and bootstrapped that statistics (N=10,000 repetitions). The confidence interval (Cl) was calculated by finding the interval containing 95% of the data (2.5 and 97.5 percentiles). As previously, we reasoned that if zero belonged to the 95% Cl, then, the means could not be deemed as being different. In contrast, if zero is found outside the Cl, then, slopes are different at p<0.05. The results of our analysis show that slopes were different between low and high stiffness conditions (Cl: 10.5 to 38.3).

In contrast, and in agreement with previous studies, grip force exhibited a first local peak that coincided with a peak of acceleration that occurred early after movement onset. Indeed, Figure 4C shows that this delay varied around zero on average for all participants and held true without exception on a subject basis (Fig. 4D). The ANOVA confirmed this observation and failed to report any effect on the lag between these two peaks (all F<0.2, all p>0.551). A t-test revealed its value was not different from zero (mean=-2.3ms; t_34_=0.3, p=0.365).

The system responds by switching the grip force lag to a different control strategy following a monotonic change in the environment. We hypothesize that this switch is triggered by a change in the value of a discrete variable that indicates the need for a qualitative change in the behavior of the system. One immediate candidate for such variable is the crossing of a stiffness threshold. Indeed, the switching in both increasing and decreasing series occurs around k=147N/m. Interestingly, while the lag was not statistically different between the two block conditions, there were two well identified global maxima in the peaks of the grip force as a function of stiffness. There is an hysteresis in the peak grip force but not the lag, across stiffness. This suggests that the motor system first accumulates some evidence about crossing some threshold and only then reacts with a switch in the feedforward control strategy to adjust the grip force peak.

Crossing a stiffness threshold is not the only possible switching variable. In previous investigations, acceleration was shown to be a key information to perform tasks involving, for instance, eye-hand coordination (White et al., 2012; Binsted and Elliott, 1999; Helsen et al., 2000). Interestingly, we found that the stiffness threshold that we identified previously marked the transition between positive and negative accelerations at the time of force onset, that is, whether the hand of the participant was accelerating or decelerating when forces started acting on the hand. We identified, for each participant, the switch in the acceleration sign at impact, and plotted the switch in strategy as a function of the switch in the sign of acceleration at impact. The correlation was statistically significant across participants (r=0.36, p=0.030). This highlights that a correlation exists at a subject-level between these two variables, which however, does not imply causation.

## Discussion

We set out to understand the phase transition between two distinct grip force control strategies during tool-mediated interaction with elastic force fields. Participants interacted with springs with increasing or decreasing stiffness between two values and controlled grip force according to the expected dynamics in all trials, including in zero-stiffness catch trials. Peak grip force reached a maximum for an average stiffness of 147N/m. Participants exhibited different qualitative behaviors related to the lag between peak grip force and elastic force in function of stiffness; for most of them, the lag increased with stiffness upon a certain threshold. Based on these observations, we suggest that the central nervous system acts as a hybrid controller that is characterized by continuous and discrete states, and operates a phase transition upon a specific stiffness value, potentially triggered by the stiffness value, the sign of the acceleration at the time of the initial contact with the elastic force field, or other candidate switching variables.

Some tasks are quintessentially complex, nonlinear and high dimensional, leading to postural instabilities and task uncertainties such as when we make contacts between two objects. Our results support a view according to which the central nervous system switches strategy in grip force control in the face of locally unstable tasks. Participants unconsciously modulate the lag between peak grip force and (expected, in catch trials) peak elastic load force. Stiff trials produce larger uncontrollable transitory forces (see Fig. 6 in White et al., 2011). Our data is in line with the idea that long latencies are better suited to trials for which instability is largest. Indeed, as we showed previously, long latencies allow the viscoelastic properties of the skin to dissipate more energy than short latencies, for which the hand is stiffer (White et al., 2011). In other words, the latency is proportional to instability. Consistently, it was suggested that increasing the delay in a control loop may, in some cases and for certain values, improve stability (Malakhovski and Mirkin, 2006). Consequently, in stiff trials, grip force is smaller at the time when perturbations are maximal than a few tens of millisecond after. In addition to the ability to dissipate energy, lowering the forces has two other positive effects. First, the perturbing forces attributable to signal-dependent noise also decrease with lower forces (Jones et al., 2002; Hamilton et al., 2004). Second, excessive co-activation is energy greedy (Sih and Stuhmiller, 2003; Foley and Meyer, 1993). These grip force adjustment differences are happening within the range of grip forces that protect the participant from object slippage, as evidenced by the fact that none of the participants ever lost grip of the object. This suggests these grip force variations are related to additional aspects of manipulated object stabilization.

This latency was not constant. Prior studies observed this latency without attempting to experimentally control it (Serrien et al., 1999; Turrell et al., 1999; Johansson and Westling, 1988; Bleyenheuft et al., 2009; Johansson et al., 1992), and found values consistent with the maximal latency (75ms) we observed in stiff interactions between a hand-held object and a surface (White et al., 2011). In a recent study, we failed to alter that latency by changing stiffness of a virtual surface and direction of movement (White et al., 2011). This was likely the case because the stiff and soft targets were implemented with 1200N/m and 240N/m virtual springs, which were both above the stiffness values encountered here. However, in a different study, we observed modulation of latency during profound gravitational changes induced by parabolic flights that challenged participants by confronting them with fundamental environmental uncertainties (White et al., 2012).

Perhaps the most striking observation is that the central nervous system switched between grip force strategies around a certain individual threshold identified through three independent observations. First, it marked the average stiffness at which grip force peaked (Fig. 3B). Second, the piecewise linear fit had a remarkable point close to this stiffness (Fig. 4A). Third, hand acceleration at force onset reversed its sign around that threshold. It is also worth reporting that a few hundreds of milliseconds before impact, in the very same trial, grip force exhibited a local peak that was synchronized with the small yet significant load force maximum due to inertia (Fig. 4C-D). When comparing Fig. 4A-B and Fig. 4C-D, it is very clear that participants predictively control grip force very differently when they are confronted to inertia or impacts. A more subtle adjustments also hold for low and high stiffness interactions.

The switching in grip force control strategy might be coupled with a general switch between controlling movement trajectories to controlling interaction forces during tool-mediated interaction with elastic objects. Previous studies suggested that stiffness (Mugge et al., 2009; Chib et al., 2006) and stiffness discontinuity crossing (Nisky et al., 2008) lead to different weighting of position and force control in manual interaction. When participants interact with low-stiffness force fields, they control kinematics, and estimate the stiffness of the elastic field based on integration of position information with sensed forces. With increasing stiffness, the reliability of stiffness estimation deteriorates in accordance with Weber’s law (Jones and Hunter, 1990). When participants interact with elastic force fields with higher stiffness (Mugge et al., 2009; Chib et al., 2006), or more frequently cross stiffness discontinuity (Nisky et al., 2008), they favor control of interaction forces rather than the control of kinematics. When this transition happens, the central nervous system might start estimating the compliance of the elastic field rather than its stiffness, resulting again in reliable estimates. Such view is consistent with our observation that peak grip forces are largest around the transition, and are smaller for very high or very low stiffness levels. It is also strikingly consistent with the threshold of around 100-200N/m in the stiffness-dependent weighting of force and position feedback (Mugge et al., 2009).

How the brain estimates stiffness is an open question. Various computational models were proposed, including: peak force divided by perceived penetration (Pressman et al., 2006, 2008, 2011), or regression of force over position or position over force data (Nisky et al., 2011, 2008, 2010). Another proposed measure is Extended Rate Hardness, a measure of the perceived hardness of a surface based on rate of force change and penetration velocity (Han and Choi, 2010). Skin deformation accompanying the probing also likely plays a role in perception of stiffness (Quek et al., 2014, Quek et al., 2015, Farajian et al., 2017). Here, we do not attempt to spill more light on this matter, but it is possible that the estimated stiffness is used in the process of the switching.

Behavioral changes upon the value reached by a variable is common in the sensorimotor system. Accurate perceptual decisions are achieved in the face of sensory noise by integrating evidence over time into a “decision variable” that triggers an action upon reaching a criterion (Smith and Ratcliff, 2004; Kelly and O’Connell, 2013). For instance, at a neuronal level, saccade initiation occurs when the discharge rate of either single neurons or a population of neurons encoding a saccade motor plan reaches a threshold level of activity, that can be modulated upon task requirements (Jantz et al., 2013). At a more behavioral level, in ball catching tasks, participants incept their actions when time to contact hit a certain value (Lee et al., 1983).

We investigated the behavioral aspects of the switching and its potential underlying switching variable. The question remains open which neural structures operate this switching. In previous studies we have shown that the left supplementary motor area (SMA) is a crucial node in the network that processes the internal representation of object dynamics (White et al., 2013) leaving this neural structure as a potential candidate to host the decision variable that controls the phase transition. We also showed that the posterior parietal cortex (PPC) is involved in the perception of stiffness (Leib et al., 2016). Other candidate areas may include the cerebellum and the insula. Several studies reported bistable states of Purkinje cells in the cerebellum that may serve as a switching trigger (Yartsev et al., 2009). Finally, the insula seems to be involved in switching between the executive control and default networks (Sridharan et al., 2008).

We should however also underline two limitations of the present study. First, we failed to explain these results within a fully coherent, average behavior. Instead, we found some idiosyncratic changes in strategy. This new question should be addressed in a follow-up experiment aiming at identifying what caused these switches. Second, our data exhibit large variability, which is inevitable when studying the interaction between mechanical interactions and physiological processes. Future investigations should improve the technical design of these experiments. Our contribution paves the way toward using switched systems theory in modeling human motor control, and opens new research questions as to the nature of the discrete state variables that drive the switching between different control strategies.

To conclude, we show here evidence that the central nervous system adopts qualitatively different grip force controls to cope with impact-like environments. Our results show that the central nervous system acts as a switching system. Our findings may have very practical implications since human-machine interfaces nowadays involve haptic feedback, but many applications of fine object manipulation are missing haptic feedback, such as robot-assisted surgery and prosthetics.

## Acknowledgments

This research was supported by the *Institut National de la Santé et de la Recherche Médicate*, the *Conseil General de Bourgogne* » and the *Fonds européen de développement régional*. RL and IN were supported by the Helmsley Charitable Trust through the Agricultural, Biological and Cognitive Robotics Initiative of Ben-Gurion University of the Negev, and the Israeli Science Foundation (grant number 823/15).

## Author Contributions Statement

O.W. and A.K. designed the experiments, collected and analyzed data; O.W., A.K., and I.N. interpreted the results, M.B. collected data and, OW, RL, CP, MB, and IN contributed to writing the manuscript, manuscript revision, read and approved the submitted version.

## Additional Information

The authors declare no competing financial interests.

## References

Asai, Y., Tasaka, Y., Nomura, K., Nomura, T., Casadio, M., and Morasso, P. (2009). A model of postural control in quiet standing: Robust compensation of delay-induced instability using intermittent activation of feedback control. PLoS One 4. doi:10.1371/journal.pone.0006169.

Augurelle, A.-S., Penta, M., White, O., and Thonnard, J.-L. (2003). The effects of a change in gravity on the dynamics of prehension. Exp. brain Res. 148, 533–40. doi:10.1007/s00221-002-1322-3.

Ben-ltzhak, S., and Karniel, A. (2008). Minimum acceleration criterion with constraints implies bang-bang control as an underlying principle for optimal trajectories of arm reaching movements. Neural Comput. 20, 779–812. doi:10.1162/neco.2007.12-05-077.

Binsted, G., and Elliott, D. (1999). Ocular perturbations and retinal / extraretinal information: the coordination of saccadic and manual movements. 193–206.

Bleyenheuft, Y., Lefèvre, P., and Thonnard, J.-L. (2009). Predictive mechanisms control grip force after impact in self-triggered perturbations. J Mot. Behav. 41, 411–417. doi:10.3200/35-08-084.

Bottaro, A., Casadio, M., Morasso, P. G., and Sanguineti, V. (2005). Body sway during quiet standing: Is it the residual chattering of an intermittent stabilization process? Hum. Mov. Sci. 24, 588–615. doi:10.1016/j.humov.2005.07.006.

Chib, V. S., Patton, J. L., Lynch, K. M., and Mussa-lvald, F. a (2006). Haptic identification of surfaces as fields of force. J. Neurophysiol. 95, 1068–77. doi:10.1152/jn.00610.2005.

Craik, K. J. W. (1947). Theory of the human operator in control systems. I. the operator as an engineering system. Br. J. Psychol. 38, 56–61. doi:10.1111/j.2044-8295.1947.tb01141.x.

Danion, F., and Sarlegna, F. R. (2007). Can the human brain predict the consequences of arm movement corrections when transporting an object? Hints from grip force adjustments. J. Neurosci. 27, 12839–43. doi:10.1523/JN EU ROSCI.3110-07.2007.

Descoins, M., Danion, F., Bootsma, R. J., and Fre, D.Æ. (2006). Predictive control of grip force when moving object with an elastic load applied on the arm. Exp. Brain Res. 172, 331–342. doi:10.1007/s00221-005-0340-3.

Doeringer, J. A., and Hogan, N. (1998). Intermittency in Preplanned Elbow Movements Persists in the Absence of Visual Feedback. J. Neurophysiol. 80, 1787–1799. Available at: http://jn.physiology.org/content/80/4/1787.abstract%5Cnpapers2://publication/uuid/657071FB-AFFF-4B57-B7FE-0D3D6B742C97.

Ebner-karestinos, D., Thonnard, J., and Bleyenheuft, Y. (2016). Precision Grip Control while Walking Down a Stair Step. PLoS One, 1–13. doi:10.5061/dryad.kf4tr.

Farajian M, Leib R, Zaidenberg T, Mussa-Ivaldi F, Nisky I (2017) Stretching the skin of the fingertip creates a perceptual and motor illusion of touching a harder spring. bioRxiv 203604. https://doi.org/10.1101/203604

Flanagan, J. R., Tresilian, J., and Wing, A. M. (1993). Coupling of grip force and load force during arm movements with grasped objects. Neurosci. Lett. 152, 53–56. doi:10.1016/0304-3940(93)90481-Y.

Flanagan, J. R., Vetter, P., Johansson, R. S., Wolpert, D. M., and Kl, O. (2003). Prediction Precedes Control in Motor Learning. Curr. Biol. 13,146-150.

Flanagan, J. R., and Wing, A. M. (1995). The stability of precision grip forces during cyclic arm movements with a hand-held load. Exp. Brain Res. 105, 455–464. doi:10.1007/BF00233045.

Foley, J. M., and Meyer, R. a (1993). Energy cost of twitch and tetanic contractions of rat muscle estimated in situ by gated 31P NMR. NMR Biomed. 6, 32–38.

Gawthrop, P., Lee, K. Y., Halaki, M., and O’Dwyer, N. (2013). Human stick balancing: An intermittent control explanation. Biol. Cybern. 107, 637–652. doi:10.1007/s00422-013-0564-4.

Gawthrop, P., Loram, I., Gollee, H., and Lakie, M. (2014). Intermittent control models of human standing: Similarities and differences. Biol. Cybern. 108, 159–168. doi:10.1007/s00422-014-0587-5.

Gross, J., Timmermann, L, Kujala, J., Dirks, M., Schmitz, F., Salmelin, R., and Schnitzler, a (2002). The neural basis of intermittent motor control in humans. Proc. Natl. Acad. Sci. U. S. A. 99, 2299–2302. doi:10.1073/pnas.032682099.

Gysin, P., Kaminski, T. R., Hass, C. E., Grobet, C. E., and Gordon, A. M. (2008). Effects of gait variations on grip force coordination during object transport. J. Neurophysiol. 100, 2477–85. doi:10.1152/jn.90561.2008.

Hamilton, A. F. de C., Jones, K. E., and Wolpert, D. M. (2004). The scaling of motor noise with muscle strength and motor unit number in humans. Exp. Brain Res. 157, 417–430. doi:10.1007/s00221-004-1856-7.

Han, G., and Choi, S. (2010). Extended rate-hardness: A measure for perceived hardness, in Lecture Notes in Computer Science (including subseries Lecture Notes in Artificial Intelligence and Lecture Notes in Bioinformatics), 117–124. doi:10.1007/978-3-642-14064-8_18.

Helsen, W. F., Elliott, D., Starkes, J. L, and Ricker, K. L. (2000). Coupling of eye, finger, elbow, and shoulder movements during manual aiming. J. Mot. Behav. 32, 241–248. doi:10.1080/00222890009601375.

Jantz, J. J., Watanabe, M., Everling, S., and Munoz, D. P. (2013). Threshold mechanism for saccade initiation in frontal eye field and superior colliculus. J. Neurophysiol. 109, 2767–80. doi:10.1152/jn.00611.2012.

Johansson, R. S., Riso, R., Häger, C., and Bäckström, L. (1992). Somatosensory control of precision grip during unpredictable pulling loads -1. Changes in load force amplitude. Exp. Brain Res. 89, 181–191. doi:10.1007/BF00229015.

Johansson, R. S., and Westling, G. (1988). Programmed and triggered actions to rapid load changes during precision grip. Exp. brain Res. 71, 72–86. Available at: http://www.ncbi.nlm.nih.gov/pubmed/3416959.

Johansson, R. S., and Westling, G. (1984). Roles of glabrous skin receptors and sensorimotor memory in automatic control of precision grip when lifting rougher or more slippery objects. Exp. Brain Res. 56, 550–564. doi:10.1007/BF00237997.

Jones, K. E., Hamilton, A. F., and Wolpert, D. M. (2002). Sources of signal-dependent noise during isometric force production. J. Neurophysiol. 88, 1533–44. Available at: http://www.ncbi.nlm.nih.gov/pubmed/12205173.

Jones, L. A., and Hunter, I. W. (1990). A perceptual analysis of stiffness. Exp. brain Res. 79, 150–156.

Karniel, A. (2011). Open Questions in Computational Motor Control. J. Integr. Neurosci. 10, 385–411. doi:10.1142/S0219635211002749.

Kelly, S. P., and O’Connell, R. G. (2013). Internal and external influences on the rate of sensory evidence accumulation in the human brain. J. Neurosci. 33, 19434–41. doi:10.1523/JNEUROSCI.3355-13.2013.

Kelso, J. a. S. (1984). Phase transitions and critical behavior in human bimanual coordination. Am. J. Physiol. 246, R1000–4. Available at: http://www.ncbi.nlm.nih.gov/pubmed/6742155.

Lee, D. N., Young, D.S., Reddish, P.E., Lough, S., and Clayton, T. M. (1983). Visual timing in hitting an accelerating ball. Q. J. Exp. Psychol. A. 35, 333–346. doi: 10.1080/14640748308402138.

Leib, R., Karniel, A., and Nisky, I. (2015). The effect of force feedback delay on stiffness perception and grip force modulation during tool-mediated interaction with elastic force fields. J. Neurophysiol. 113, 3076–3089. Available at: http://jn.physiology.org/content/early/2015/02/19/jn.00229.2014.abstract.

Leib, R., and Karniel, a. (2012). Minimum acceleration with constraints of center of mass: a unified model for arm movements and object manipulation. J. Neurophysiol. 108, 1646–1655. doi:10.1152/jn.00224.2012.

Leib, R., Mawase, F., Karniel, A., Donchin, O., Rothwell, J., Nisky, I., and Davare, M. (2016). Stimulation of PPC Affects the Mapping between Motion and Force Signals for Stiffness Perception But Not Motion Control. J. Neurosci. 36, 10545–10559. doi:10.1523/JNEUROSCI.1178-16.2016.

Levy-Tzedek, S., Krebs, H.I., Song, D., Hogan, N., and Poizner, H. (2010). Non-monotonicity on a spatio-temporally defined cyclic task: Evidence of two movement types? Exp. Brain Res. 202, 733–746. doi:10.1007/s00221-010-2176-8.

Levy-Tzedek, S., Ben Tov, M., and Karniel, A. (2011). Early switching between movement types: Indication of predictive control? Brain Res. Bull. 85, 283–288. doi:10.1016/j.brainresbull.2010.11.010.

Liberzon, D. (2003). Basic Concepts. In: Switching in Systems and Control. Boston MA. Birkhäuser Boston.

Lin, H., and Antsaklis, P. J. (2009). Stability and stabilizability of switched linear systems: A survey of recent results. IEEE Trans. Automat. Contr. 54, 308–322. doi:10.1109/TAC.2008.2012009.

Loewenstein, Y., Mahon, S., Chadderton, P., Kitamura, K., Sompolinsky, H., Yarom, Y., and Häusser, M. (2005). Bistability of cerebellar Purkinje cells modulated by sensory stimulation. Nat. Neurosci. 8, 202–11. doi:10.1038/nnl393.

Loram, I. D., Gollee, H., Lakie, M., and Gawthrop, P. J. (2011). Human control of an inverted pendulum: is continuous control necessary? Is intermittent control effective? Is intermittent control physiological? J. Physiol. 589, 307–324. doi:10.1113/jphysiol.2010.194712.

Malakhovski, E., and Mirkin, L. (2006). On stability of second-order quasi-polynomials with a single delay. Automatica 42, 1041–1047. doi:10.1016/j.automatica.2006.01.024.

Margaliot, M., and Liberzon, D. (2006). Lie-algebraic stability conditions for nonlinear switched systems and differential inclusions. Syst. Control Lett. 55, 8–16. doi:10.1016/j.sysconle.2005.04.011.

Miall, R. C., Weir, D.J., and Stein, J. F. (1993). Intermittency in human manual tracking tasks. J. Mot. Behav. 25, 53–63. doi: 10.1080/00222895.1993.9941639.

Mugge, W., Schuurmans, J., Schouten, A.C., and van der Helm, F. C. T. (2009). Sensory weighting of force and position feedback in human motor control tasks. J. Neurosci. 29, 5476–82. doi:10.1523/JNEUROSCI.OH6-09.2009.

Navas, F., and Stark, L. (1968). Sampling or intermittency in hand control system dynamics. Biophys. J. 8, 252–302. doi: 10.1016/S0006-3495(68)86488-4.

Neilson, P. D., Neilson, M.D., and O’Dwyer, N. J. (1988). Internal models and intermittency: A theoretical account of human tracking behavior. Biol. Cybern. 58, 101–112. doi:10.1007/BF00364156.

Nisky, I., Baraduc, P., and Karniel, A. (2010). Proximodistal gradient in the perception of delayed stiffness. J. Neurophysiol. 103, 3017–26. doi:10.1152/jn.00939.2009.

Nisky, I., Mussa-Ivaldi, F. A., and Karniel, A. (2008). A regression and boundary-crossing-based model for the perception of delayed stiffness. IEEE Trans. Haptics 1, 73–83. doi:10.1109/TOH.2008.17.

Nisky, I., Pressman, A., Pugh, C.M., Mussa-Ivaldi, F. A., and Karniel, A. (2011). Perception and action in teleoperated needle insertion. IEEE Trans. Haptics 4, 155–166. doi:10.1109/TOH.2011.30.

Nowak, D. A., Hermsdörfer, J., Schneider, E., and Glasauer, S. (2004). Movin ag objects in a rotating environment: rapid prediction of Coriolis and centrifugal force perturbations. Exp. brain Res. 157, 241–54. doi:10.1007/s00221-004-1839-8.

Pressman, A., Karniel, A., and Mussa-Ivaldi, F. A. (2011). How soft is that pillow? The perceptual localization of the hand and the haptic assessment of contact rigidity. J. Neurosci. 31, 6595–6604. doi:10.1523/JN EU ROSCI.4656-10.2011.

Pressman, A., Karniel, A., and Mussa-Ivaldi, F. A. (2006). Perception of delayed stiffness, in Proceedings of the First IEEE/RAS-EMBS International Conference on Biomedical Robotics and Biomechatronics, 2006, BioRob 2006, 905–910. doi:10.1109/BIOROB.2006.1639206.

Pressman, A., Nisky, I., Karniel, A., and Mussa-Ivaldi, F. a. (2008). Probing Virtual Boundaries and the Perception of Delayed Stiffness. Adv. Robot. 22, 119–140. doi:10.1163/156855308X291863.

Quek Z, Schorr S, Nisky I, Provancher W, Okamura A (2015) Sensory Substitution and Augmentation Using 3-Degree-of-Freedom Skin Deformation. IEEE Transactions on Haptics 8:209–221. doi: 10.1109/TOH.2015.2398448

Quek ZF, Schorr SB, Nisky I, Okamura AM, Provancher WR (2014) Augmentation Of Stiffness Perception With a 1-Degree-of-Freedom Skin Stretch Device. IEEE Transactions on Human-Machine Systems, 44:731–742. doi: 10.1109/TH MS.2014.2348865

Sarlegna, F. R., Baud-Bovy, G., and Danion, F. (2010). Delayed visual feedback affects both manual tracking and grip force control when transporting a handheld object. J. Neurophysiol. 104, 641–653. doi:10.1152/jn.00174.2010.

Serrien, D. J., Kaluzny, P., Wicki, U., and Wiesendanger, M. (1999). Grip force adjustments induced by predictable load perturbations during a manipulative task. Exp. brain Res. 124, 100–6. Available at: http://www.ncbi.nlm.nih.gov/pubmed/9928794.

Sih, B. L., and Stuhmiller, J. H. (2003). The metabolic cost of force generation. Med. Sci. Sports Exerc. 35, 623–629. doi:10.1249/01.MSS.0000058435.67376.49.

Smith, P. L., and Ratcliff, R. (2004). Psychology and neurobiology of simple decisions. Trends Neurosci. 27, 161–168. doi:10.1016/j.tins.2004.01.006.

Squeri, V., Masia, L., Casadio, M., Morasso, P., and Vergaro, E. (2010). Force-Field compensation in a manual tracking task. PLoS One 5. doi:10.1371/journal.pone.0011189.

Sridharan, D., Levitin, D.J., and Menon, V. (2008). A critical role for the right fronto-insular cortex in switching between central-executive and default-mode networks. Proc Natl Acad Sci USA 105, 12569–12574. doi:10.1073/pnas.0800005105.

Turrell, Y. N., Li, F.X., and Wing, A. M. (1999). Grip force dynamics in the approach to a collision, in Experimental Brain Research, 86–91. doi:10.1007/s002210050822.

White, O. (2015). The brain adjusts grip forces differently according to gravity and inertiaffl: a parabolic flight experiment. Front. Integr. Neurosci. 9,1–10. doi:10.3389/fnint.2015.00007.

White, O., Davare, M., Andres, M., and Olivier, E. (2013). The role of left supplementary motor area in grip force scaling. PLoS One 8. doi:10.1371/journal.pone.0083812.

White, O., Dowling, N., Bracewell, R.M., and Diedrichsen, J. (2008). Hand interactions in rapid grip force adjustments are independent of object dynamics. J. Neurophysiol. 100, 2738–45. doi:10.1152/jn.90593.2008.

White, O., Lefèvre, P., Wing, A.M., Bracewell, R.M., and Thonnard, J. L. (2012). Active Collisions in Altered Gravity Reveal Eye-Hand Coordination Strategies. PLoS One 7. doi:10.1371/journal.pone.0044291.

White, O., Thonnard, J.-L., Wing, a M., Bracewell, R.M., Diedrichsen, J., and Lefèvre, P. (2011). Grip force regulates hand impedance to optimize object stability in high impact loads. Neuroscience 189, 269–76. doi:10.1016/j.neuroscience.2011.04.055.

Wicks, M., Peleties, P., and DeCarlo, R. (1998). Switched controller synthesis for the quadratic stabilisation of a pair of unstable linear systems. Eur. J. Control 4, 140–147. doi:10.1016/S0947-3580(98)70108-6.

Yartsev, M. M., Givon-Mayo, R., Maller, M., and Donchin, O. (2009). Pausing purkinje cells in the cerebellum of the awake cat. Front. Syst. Neurosci. 3, 2. doi:10.3389/neuro.06.002.2009.

